# Testing the efficacy of lionfish traps in the northern Gulf of Mexico

**DOI:** 10.1101/2020.03.16.993543

**Authors:** Holden E. Harris, Alexander Q. Fogg, Stephen R. Gittings, Robert N. M. Ahrens, Micheal S. Allen, William F. Patterson

## Abstract

Spearfishing is currently the primary approach for removing invasive Indo-Pacific lionfish *(Pterois volitans/miles* complex) to mitigate their impacts to western Atlantic marine ecosystems. However, a substantial portion of lionfish spawning biomass is beyond the depth limits of SCUBA. Innovative technologies may offer an alternative means to target lionfish and allow for the development of a deepwater lionfish trap fishery, but the removal efficiency and potential environmental impacts of traps have not been evaluated. We tested a collapsible non-containment trap (the ‘Gittings trap’) near artificial reefs in the northern Gulf of Mexico. A total of 327 lionfish and 28 native fishes (4 of which regulated species) recruited to traps (i.e., number of fish observed within the trap footprint) during 82 trap sets, catching 144 lionfish and 29 native species. Lionfish recruitment was highest for single (versus paired) traps deployed <15 m from reefs with a 1-day soak time, for which mean lionfish and native species recruitment per trap were approximately 5 and 0.1, respectively. Gittings traps selected for larger lionfish (mean size 277 mm total length) compared to spearfishing (mean size 258 mm). Community impacts from Gittings traps appeared minimal given recruitment patterns and catch rates were >10X higher for lionfish versus native fishes. Gittings traps did not move on the bottom during two major storm events; however, further testing will be necessary to test trap movement with surface buoys. Additional research should also focus on design and operational modifications to improve Gittings trap deployment success (68% successfully opened on the seabed) and reduce lionfish escapement (56% escaped from traps upon retrieval). While removal efficiency for lionfish (12–24%) was far below that of spearfishing, study results demonstrate Gittings traps are suitable for future testing on deepwater natural reefs, which constitute >90% of the region’s reef habitat.

## Introduction

Invasive Indo-Pacific lionfish (*Pterois volitans/miles* complex, hereafter “lionfish”) are now well established in the western Atlantic, including the Caribbean Sea and Gulf of Mexico [1] and have recently invaded the Mediterranean Sea [2]. Lionfish occupy a wide diversity of invaded marine habitats, including coral reefs, subtropical artificial and natural reefs [3], seagrass beds [4], mangroves [5], estuaries [6], mesophotic reefs [7–9], and upper continental slope reefs [10]. High population densities of lionfish [3,11] have caused reductions in native reef fish abundances [12,13], altered marine communities [14,15], and likely exacerbate current stressors on marine systems [16,17]. As invasive lionfish populations do not appear to be controlled by native predators [18–20], reducing lionfish biomass a top priority for marine resource managers [21,22].

The current capacity to remove lionfish is primarily by spearfishing [21,23]. However, lionfish have been observed at depths >300 m [10] and spearfishing removals for lionfish are limited to SCUBA accessible depths (<40 m). Over 557,000 km^2^ of benthic habitat in western Atlantic invaded range of lionfish lies within mesophotic depths of 40–300 m [24] where lionfish populations densities are often higher than shallower depths [9,25–27]. Deepwater lionfish populations likely disrupt food webs on mesophotic reefs [7] and provide refuge for larger and more fecund individuals [28]. These protected source populations can provide larvae for sink regions [29,30] and undermine shallow-water control efforts [8,28]. Population and ecosystem models predict high levels of fishing mortality over a broad geographic range will be necessary to control lionfish populations on a regional scale [15,29,31,32].

Innovative harvest technologies have been proposed for deepwater lionfish removals, including modifications to existing spiny lobster traps [33], weaponized remotely operated vehicles [22,34], and novel trap designs [35]. Collaborative work by members of non-profit organizations, Florida Fish and Wildlife Conservation Commission and US National Oceanic and Atmospheric Administration have resulted in lionfish trap prototypes [36], which have been further developed into a non-containment trap model [37]. The collapsible ‘Gittings trap’ is designed to allow lionfish and other marine species to freely move over the trap’s footprint with traps closing during retrieval (**Table 1**). Gittings traps are made from common and inexpensive materials, allowing for construction on small islands where specialized materials may be difficult to source. Ultimately, a low cost of production could expand fishing capacity for the nascent lionfish market, which is currently constrained by inconsistent supply [38,39]. A commercial deepwater lionfish fishery may further offer alternative livelihood strategies for fishermen and improve coastal food security [40]. However, it will be critical to further evaluate this new harvest gear for potential undesirable effects (e.g., bycatch, habitat damage, entanglement, or ghost fishing) before it is permitted and widely implemented. Traditional fish traps have a broad catch composition [41] and their widespread use has contributed to severe overfishing on many Caribbean coral reef systems [41,42]. Given the potential for bycatch and overfishing, moratoriums on fish traps have been in place in US Atlantic and Gulf of Mexico waters for decades [43], with the exception of a limited trap fishery for Atlantic black sea bass, *Centropristis striata* [44].

**Table 1.**
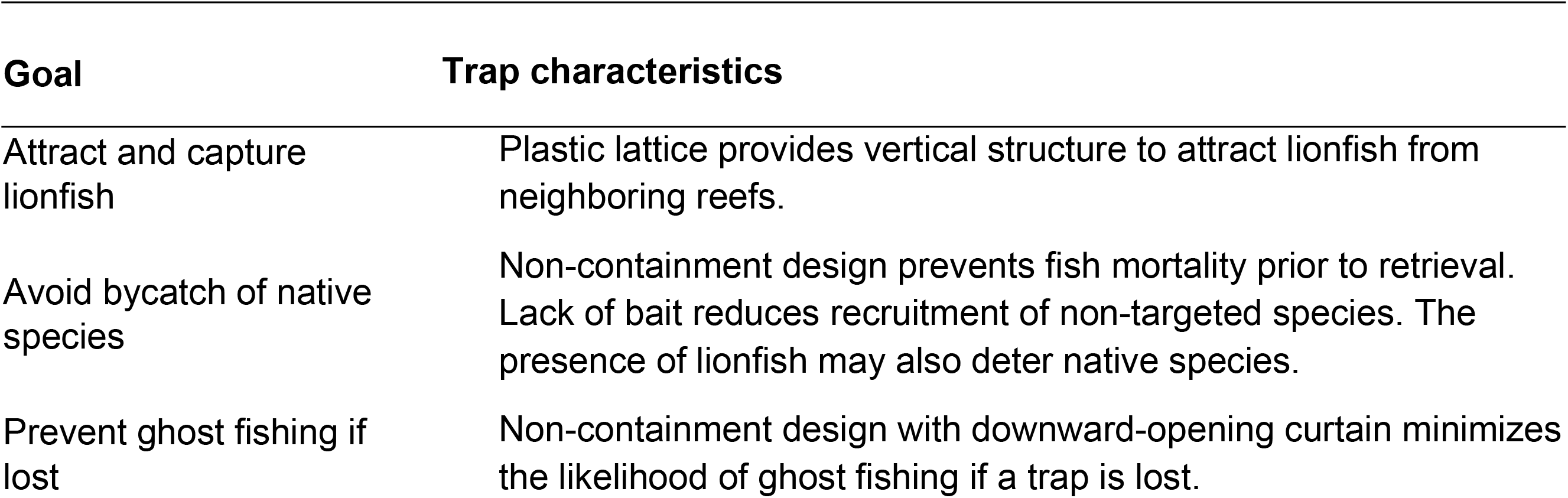
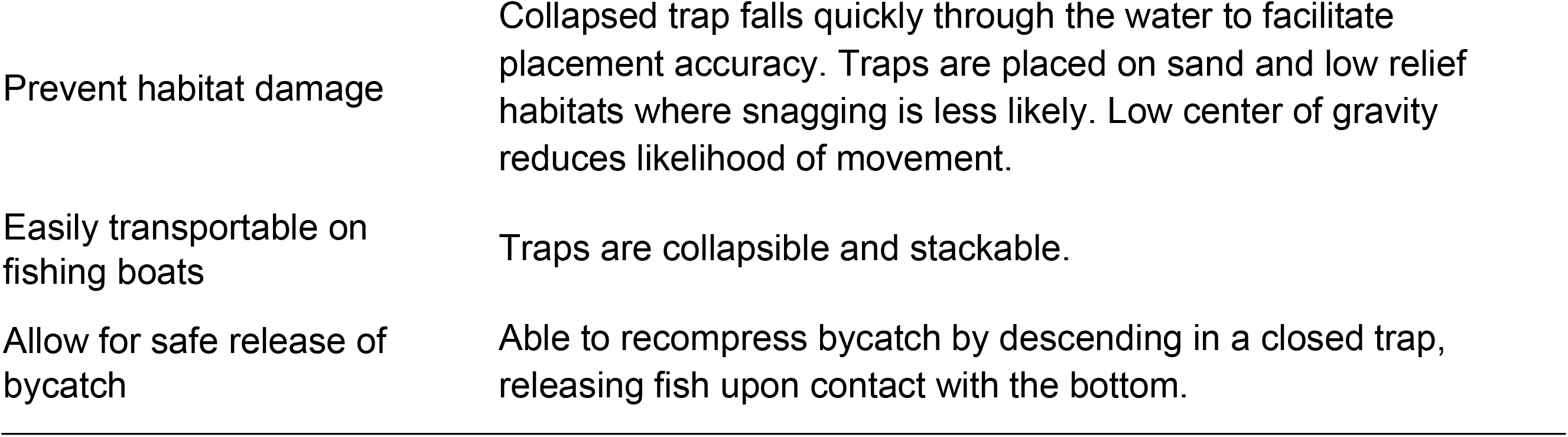
Gittings trap characteristics. Goals of the Gittings trap with design attributes employed to achieve those goals.

| Goal | Trap characteristics |
| --- | --- |
| Attract and capture lionfish | Plastic lattice provides vertical structure to attract lionfish from neighboring reefs. |
| Avoid bycatch of native species | Non-containment design prevents fish mortality prior to retrieval. Lack of bait reduces recruitment of non-targeted species. The presence of lionfish may also deter native species. |
| Prevent ghost fishing if lost | Non-containment design with downward-opening curtain minimizes the likelihood of ghost fishing if a trap is lost. |
| Prevent habitat damage | Collapsed trap falls quickly through the water to facilitate placement accuracy. Traps are placed on sand and low relief habitats where snagging is less likely. Low center of gravity reduces likelihood of movement. |
| Easily transportable on fishing boats | Traps are collapsible and stackable. |
| Allow for safe release of bycatch | Able to recompress bycatch by descending in a closed trap, releasing fish upon contact with the bottom. |

Here, we report results from testing Gittings traps near artificial reefs in the northern Gulf of Mexico (nGOM). We examined gear performance and how lionfish and native species recruitment (i.e., number of fish observed within the trap footprint) and catches (i.e., fish landed aboard the vessel) were affected by the number of Gittings traps deployed, their distance from the artificial reef, soak time, and lionfish density on the adjacent reef. Lionfish size was compared between Gittings trap catches and *in situ* size distributions. Gear testing Gittings trap performance involved evaluating deployment success (% of traps that successfully opened on the seabed), lionfish escapement (% of individuals that escaped traps upon retrieval), and whether traps moved on the seabed while deployed. We consider how findings from this initial study can inform further research and development of lionfish traps and innovative harvest technologies to control deepwater lionfish populations.

## Methods

Twelve Gittings traps were constructed in May and June 2018. Gittings traps have hinged jaws that allow for the trap to remain closed and travel vertically through the water column (**Fig 1A**) then open on the bottom (**Fig 1B**). Trap jaws were made from 4.5 m long sections of #6 rebar (19 mm diameter). Jaws were bent into two half-hoops with a curved extension on one end of each jaw to act as deflectors for opening the trap. The jaws pivot around a 2 m long center axle made with #6 round bar. The axle and jaws are connected with two metal hinge plates (4 × 10 cm) each with holes approximately 21 mm in diameter. Trap netting consisted of 3 m^2^ of 22 mm diameter mesh nylon netting (#420 green knotless). A sheet of plastic lattice (71 cm × 75 cm with 2.5 cm openings) provided vertical structure for attracting lionfish (**Fig. 1C**). A two-line harness was attached to the apex of each trap jaw using 12-strand Dyneema fiber rope (Amsteel Blue, 2.78 mm diameter). To prevent this line from fouling within the trap, an inline syntactic foam float was secured at the apex of the harness. An instructional video for building similar Gittings traps is provided at https://youtu.be/ta8WInxyXFA.

**Fig 1.**
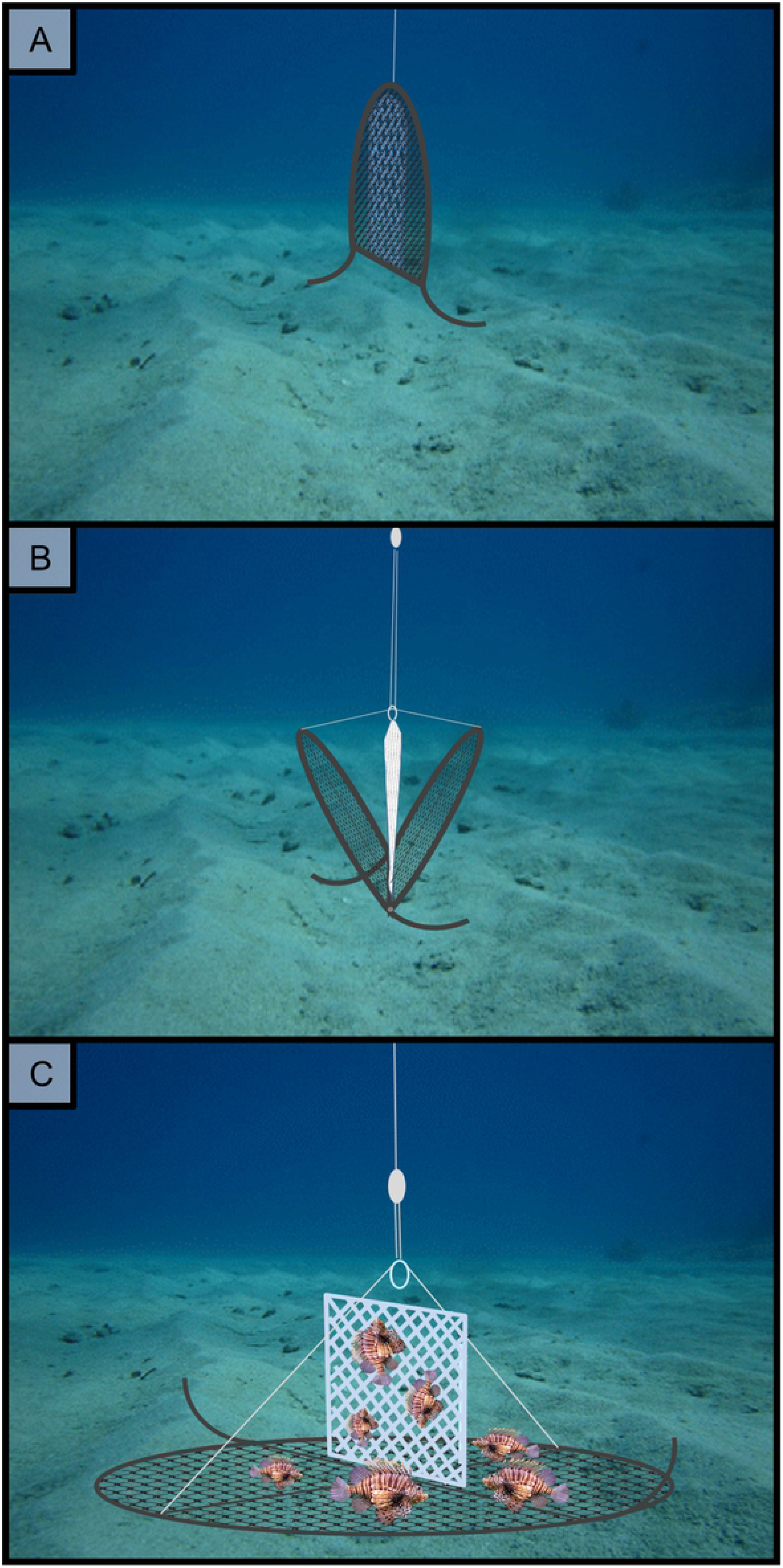
Schematic of Gittings lionfish trap deployment. A) Traps are designed to descend closed and B) open when the deflectors contact the seafloor. C) The traps remain open during deployment, then close when the trap is ascended during retrieval.

Gittings traps were deployed at depths of 33–37 m near eight artificial reefs on the nGOM Florida shelf. Reefs included four poultry transport units (i.e., chicken coops), one steel pyramid, one cement mixer, and two military tanks. These were located approximately 30 km south of Destin, Florida (**Fig. 2**). Lionfish density at each reef was surveyed immediately prior to trap deployment and trap retrieval. Surveys were conducted with point-counts by SCUBA divers within a 15 m wide cylinder with a given reef at the center. Lionfish abundance was estimated via diver counts of lionfish outside and within the structure of the reef [45,46]. Lionfish density (fish per 100 m^2^) was computed as abundance (number lionfish counted) divided by the circular area sampled (177 m^2^).

**Fig 2.**
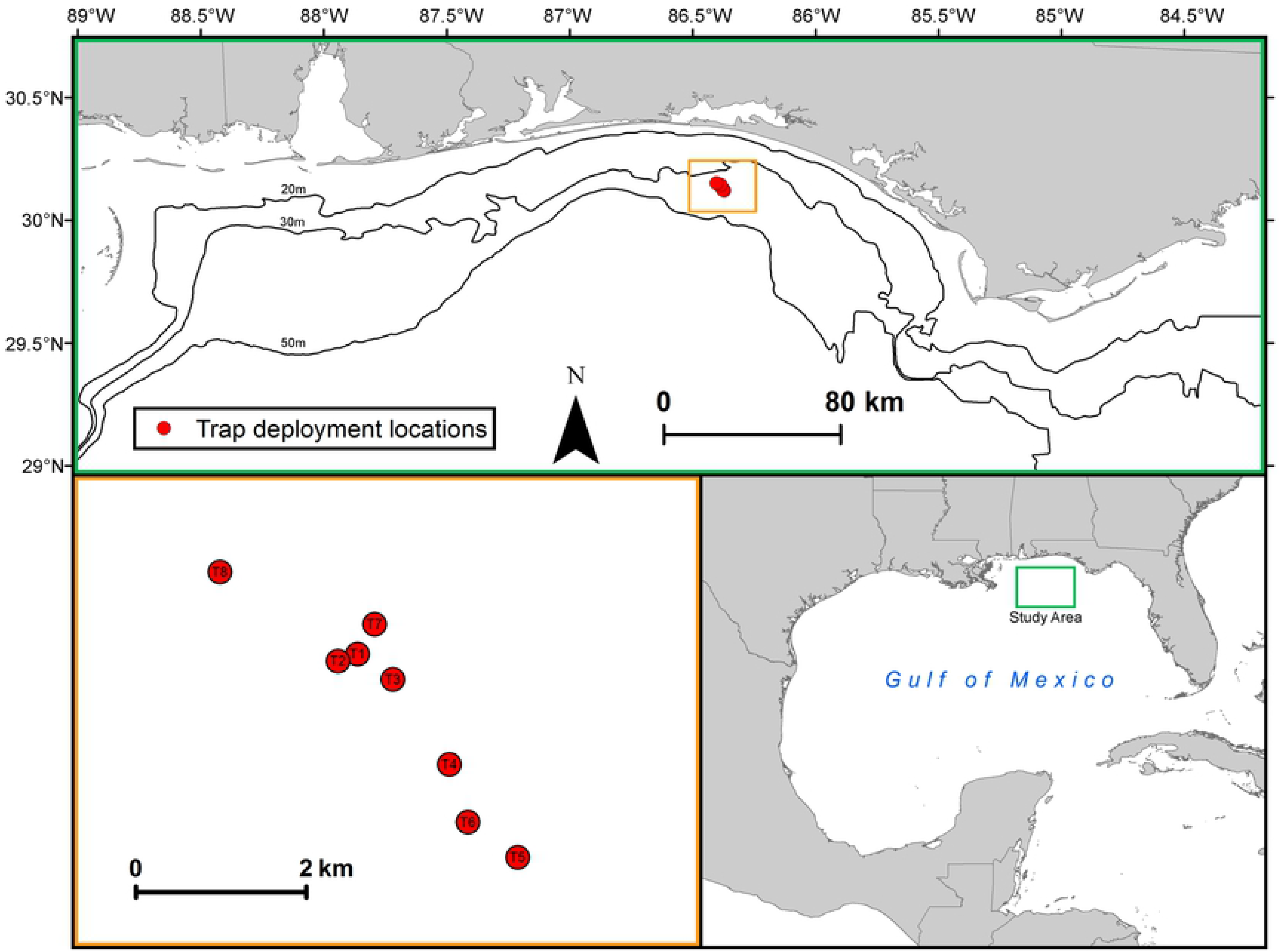
Map of study sites for testing Gittings traps. Traps were deployed in depths of 33–37 m near small artificial reefs in the northern Gulf of Mexico approximately 30 km offshore NW Florida. Deployment effects (i.e., trap array, distance to reef, and soak time) were randomized during each deployment.

Trap deployments and retrievals (n = 58) were conducted during June to December 2018. Deployment factors for number of traps (two levels: single or paired), distance to reef [three levels: near (5 m), intermediate (15 m), or far (65 m)], and soak time (categorical with five levels: 3–6 h, 1 d, 4–5 d, 8 d, or 12–14 d) were randomized for each deployment. Traps were deployed from the vessel and allowed to descend freely. Deployment success (i.e., traps landing upright and opened) for each trap deployment (n = 82) was noted by SCUBA divers during the site survey and unsuccessful deployments were corrected. Traps were retrieved by SCUBA divers using two 22-kg lift bags filled with air (video of trap retrieval provided at https://youtu.be/Tf8K6ZwQV_Y). Recruitment of lionfish and native species to the footprint of the trap was documented by a SCUBA diver prior to retrieval and subsequently compared to the numbers of each in the catch.

Generalized linear mixed models (GLMMs) were computed to test the effect of deployment factors (number of traps, distance to artificial reef, and soak time) on lionfish and native reef fish recruitment to and catch by Gittings traps (**Eq. 1**). Lionfish density on adjacent reefs was included as a covariate. Individual reefs were sampled multiple times with different deployment configurations and soak times, thus reef site was included in the GLMMs as a random effect (random intercept) and assumed to be normally distributed with a mean of zero and variance σ^2^. Deployment factors and the lionfish density covariate were evaluated at an experiment-wise error rate (α) of 0.05. Quantile-quantile (QQ) plots were examined to determine if errors were best fit with a normal, lognormal, Poisson, or negative binomial distribution. Likelihood was estimated with Laplace approximation based on GLMM fitting and inference protocols [47]. Analyses were conducted in R (version 3.6.1) using the LME4 [48] and MASS [49] packages. See supplemental material for R code and raw data. The QQ plots can be produced by running the R code.

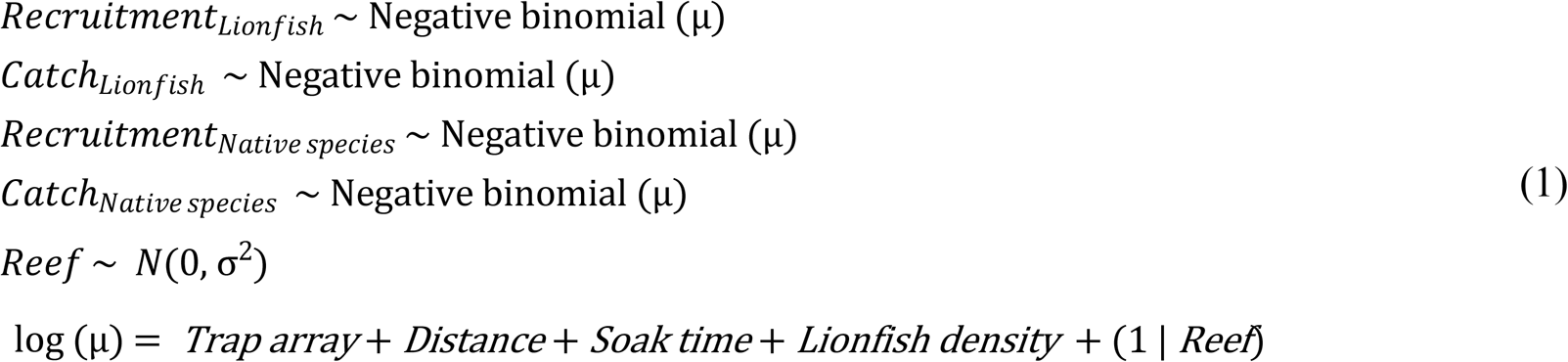

Total length (TL) was measured for all trap-caught lionfish (n = 163) and compared to that of lionfish caught via spearfishing (n = 3,063) at other artificial reefs in the study region. During spearfishing, fishers attempted to capture all lionfish observed on a given artificial reef regardless of size. A nonparametric two-sample Kolmogorov–Smirnov (KS) test was used to compare TL distributions given they were non-normal with multiple modes present. Lionfish TL data met the assumptions of the KS test, i.e., data were independent, ordinal, uncensored, ungrouped, and followed a continuous distribution.

## Results

### Gear testing

Gittings traps that were successfully deployed readily recruited lionfish from nearby artificial reefs (**Fig. 3**). Traps deployed upright and opened during 56 of 82 (68%) deployments (video of Gittings trap opening during deployment and provided at https://youtu.be/XlyNuLxEqgQ). Lionfish escapement was an issue with 56% of the lionfish that recruited to traps escaping during retrieval. Traps closed during ascent when SCUBA divers filled lift bags attached to traps with air, but this process caused traps to close slowly (~3–5 sec) allowing some lionfish to swim out of the trap before it was fully closed.

**Fig 3.**
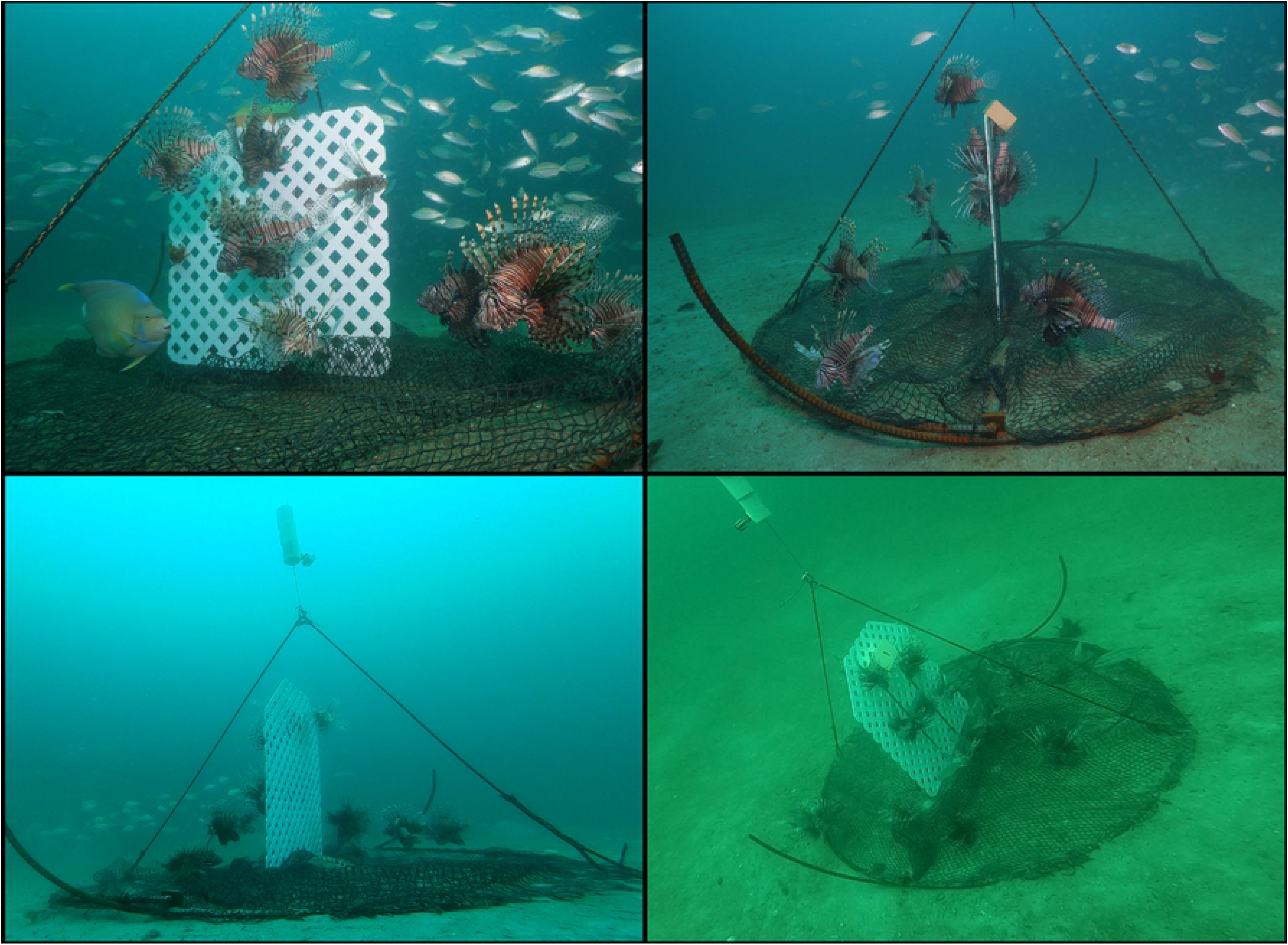
In water photos of a Gittings lionfish traps deployed near artificial reefs in the northern Gulf of Mexico. Lionfish are attracted to the structure made by piece of plastic lattice. Images: A. Fogg and H. Harris.

The potential for Gittings trap movement was tested during two severe weather events. On Sept 4–5, 2018 the center of Tropical Storm Gordon passed ~150 km west of 12 deployed traps with maximum sustained winds of 112 km/h and recorded seas >5 m. Traps were retrieved two days later, all 12 of which were upright with no change in location, although traps were heavily fouled with algae (video of trap retrieval after Tropical Storm Gordon is provided at https://youtu.be/7wZpe5fOozs). Then, on Oct 9–10, 2018 Category 5 Hurricane Michael passed ~100 km east of six deployed traps with maximum sustained winds >250 km/h and seas >15 m. Traps were unable to be retrieved for over a month due to extensive damage in the region, but upon retrieval all six traps were upright in their deployment locations. While these observations indicate high-energy storm events did not move Gittings traps on the seabed, it is currently unclear if or what amount of movement would have occurred if surface buoys had been attached to the traps.

### Lionfish and native species trap recruitment

A total of 327 lionfish recruited to traps during 82 trap sets (n = 58 deployments including paired sets) with 141 lionfish caught (**Fig. 4**). Trap bycatch of native species consisted of 28 individuals recruiting to the traps and 29 individuals caught from nine different species. One less native species was caught versus documented as recruited due to detection error by SCUBA divers. Native species catches consisted of 15 sand perch (*Diplectrum formosum)*, four tomtate (*Haemulon aurolineatum*), two bank sea bass (*Centropristis ocyurus*), two porgies (*Calamus* spp.), one soapfish (*Rypticus* spp.), and one batfish (*Ogcocephalus radiatus*). Four additional native fishes captured in traps were regulated species: two scamp (*Mycteroperca phenax*), one Gulf flounder (*Paralichthys albiguttata)*, and one blue angelfish (*Holoncanthus bermudensis*).

**Fig 4.**
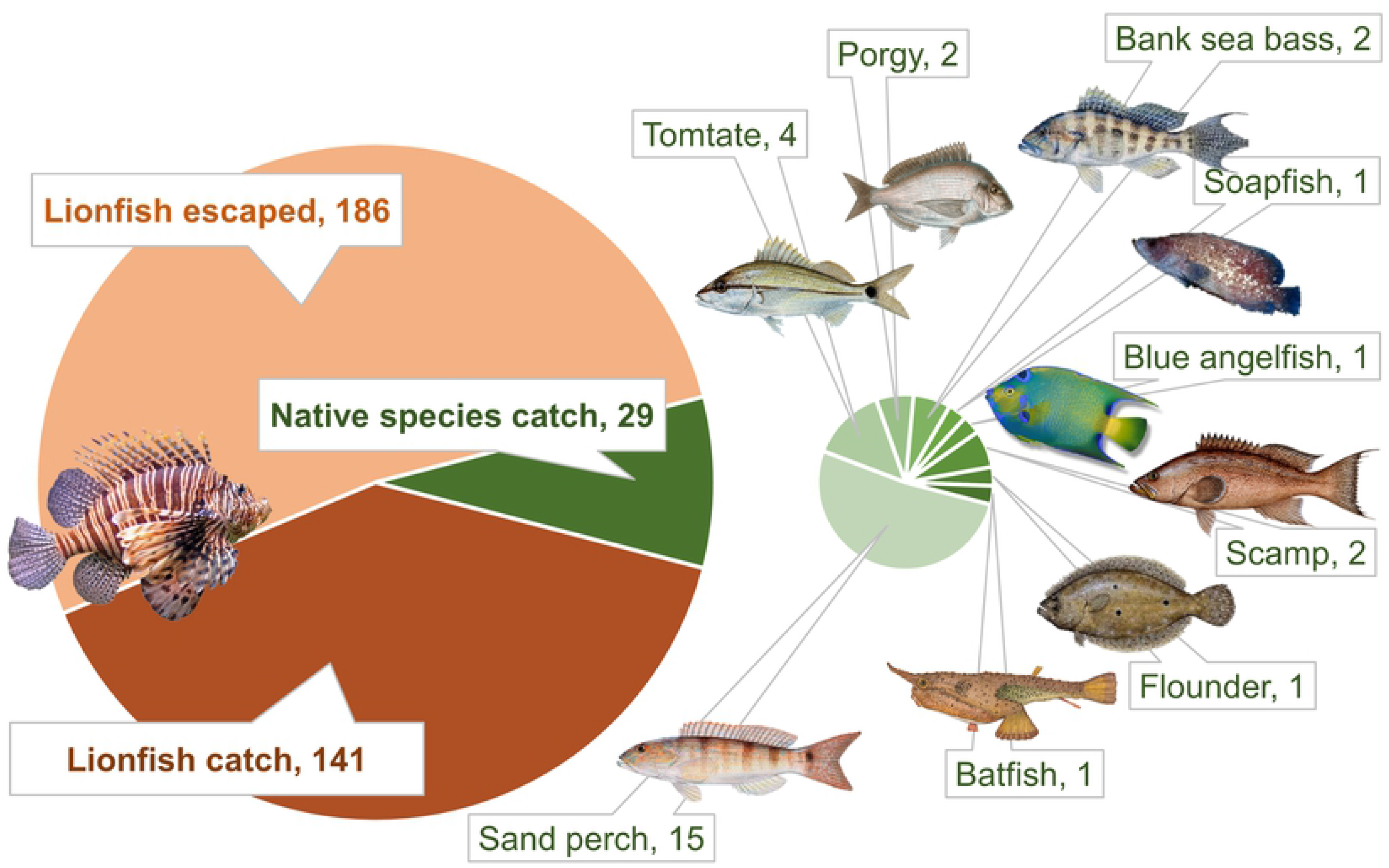
Number and species of fishes in the Gittings traps. Total counts of lionfish catch, lionfish escaped, and native species catch in traps deployed near northern Gulf of Mexico artificial reefs during 82 trap deployments. Size of pie slices correspond to proportion overall count. Fish images are not drawn to scale.

Lionfish recruitment ranged from 0 to 20 individuals and lionfish catch ranged from 0 to 14 fish per trap (video of a Gittings trap with high recruitment to is provided at https://youtu.be/1vzByPMm7hQ). QQ plots indicated the error structures for all models were best fit with a negative binomial distribution, which is typical for count data [50]. GLMM results indicated lionfish recruitment and catch were significantly affected by distance from the reef and soak time (**Table 2**). Paired traps recruited 44% fewer lionfish per trap than a single trap (**Fig. 5A**); however, the difference was not significantly different (P = 0.194) given the high variance in the data. Traps placed at close and intermediate distances (5 m and 15 m) had similar recruitment of approximately 5 lionfish per deployment (P = 0.935), while traps placed 65 m from a reef attracted almost no lionfish (P < 0.001) (**Fig. 5B**). Lionfish recruitment and catch were highest for traps deployed for 1 day and 4–5 days (**Fig. 5C**), and a soak time of 3–6 hours had significantly lower lionfish recruitment (P = 0.042). Lionfish catch was 84% lower for the longest soak time of 12–14 days (P = 0.001) while lionfish recruitment for 12–14 days was 31% lower and not statistically significant (P = 0.479), indicating escapement was the a partial driver for declines in the catch model for these trap sets. Unexpectedly, lionfish density on the adjacent artificial reefs was not a significant covariate in predicting lionfish recruitment (P = 0.152) or lionfish catch (P = 0.267) by Gittings traps.

**Table 2.**
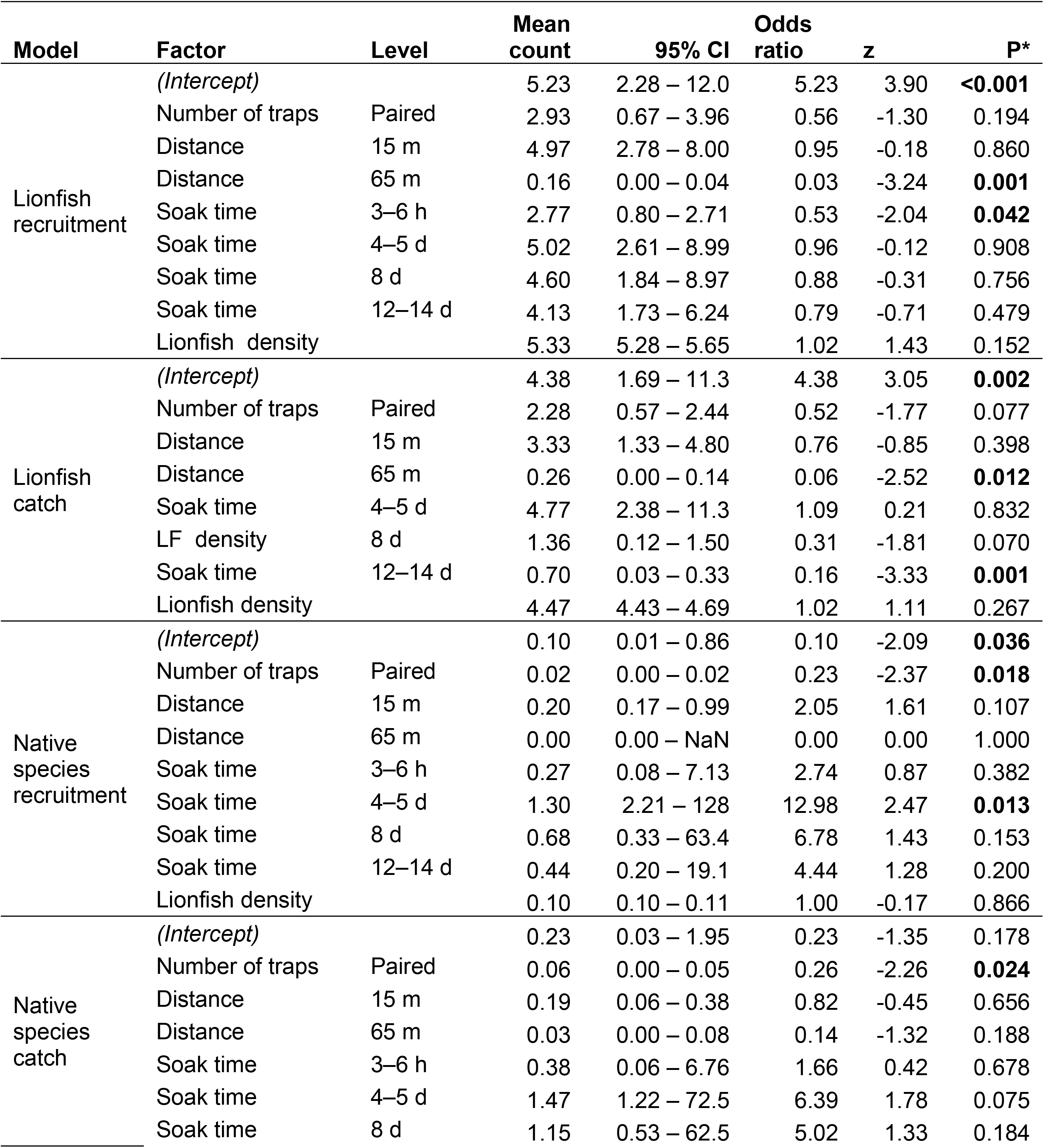
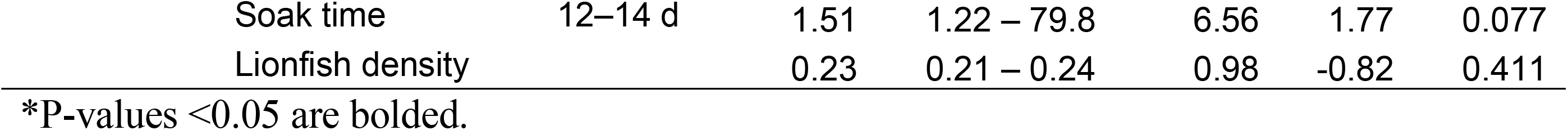
Generalized mixed model results for the effect of deployment strategies on fish recruitment and catches by Gittings traps. Effect of deployment factors and lionfish density on recruitment (fish observed within the trap footprint during retrieval) and catches (fish landed aboard the vessel) of lionfish and native reef fish species by Gittings traps tested near artificial reefs in the northern Gulf of Mexico. Factors examined included number of traps (single or paired), distance from the adjacent artificial reef (5 m, 15 m, or 65 m), soak time (3–6 hours, 1 day, 4–5 days, or 12–14 days), and source reef lionfish density (fish per 100 m^2^). Differences were tested with generalized linear mixed models (GLMM) fit with a negative binomial error distributions and reef site as a random effect. GLMM outputs show the log-linked parameter estimates for mean fish count recruited or caught per trap, associated 95% confidence intervals (CI) around mean counts, and the odds ratio (i.e., proportional effect). Odds ratio and hypothesis testing (z- and P-values) represent the difference from the GLMM intercept, i.e., the difference compared to a single trap deployed 5 m from the reef with a 1 d soak time.

| Model | Factor | Level | Mean count | 95% CI | Odds ratio | z | P* |
| --- | --- | --- | --- | --- | --- | --- | --- |
| Lionfish recruitment | (Intercept) |  | 5.23 | 2.28 – 12.0 | 5.23 | 3.90 | **<0.001** |
|  | Number of traps | Paired | 2.93 | 0.67 – 3.96 | 0.56 | -1.30 | 0.194 |
|  | Distance | 15 m | 4.97 | 2.78 – 8.00 | 0.95 | -0.18 | 0.860 |
|  | Distance | 65 m | 0.16 | 0.00 – 0.04 | 0.03 | -3.24 | **0.001** |
|  | Soak time | 3–6 h | 2.77 | 0.80 – 2.71 | 0.53 | -2.04 | **0.042** |
|  | Soak time | 4–5 d | 5.02 | 2.61 – 8.99 | 0.96 | -0.12 | 0.908 |
|  | Soak time | 8 d | 4.60 | 1.84 – 8.97 | 0.88 | -0.31 | 0.756 |
|  | Soak time | 12–14 d | 4.13 | 1.73 – 6.24 | 0.79 | -0.71 | 0.479 |
|  | Lionfish density |  | 5.33 | 5.28 – 5.65 | 1.02 | 1.43 | 0.152 |
| Lionfish catch | (Intercept) |  | 4.38 | 1.69 – 11.3 | 4.38 | 3.05 | **0.002** |
|  | Number of traps | Paired | 2.28 | 0.57 – 2.44 | 0.52 | -1.77 | 0.077 |
|  | Distance | 15 m | 3.33 | 1.33 – 4.80 | 0.76 | -0.85 | 0.398 |
|  | Distance | 65 m | 0.26 | 0.00 – 0.14 | 0.06 | -2.52 | **0.012** |
|  | Soak time | 4–5 d | 4.77 | 2.38 – 11.3 | 1.09 | 0.21 | 0.832 |
|  | LF density | 8 d | 1.36 | 0.12 – 1.50 | 0.31 | -1.81 | 0.070 |
|  | Soak time | 12–14 d | 0.70 | 0.03 – 0.33 | 0.16 | -3.33 | **0.001** |
|  | Lionfish density |  | 4.47 | 4.43 – 4.69 | 1.02 | 1.11 | 0.267 |
| Native species recruitment | (Intercept) |  | 0.10 | 0.01 – 0.86 | 0.10 | -2.09 | **0.036** |
|  | Number of traps | Paired | 0.02 | 0.00 – 0.02 | 0.23 | -2.37 | **0.018** |
|  | Distance | 15 m | 0.20 | 0.17 – 0.99 | 2.05 | 1.61 | 0.107 |
|  | Distance | 65 m | 0.00 | 0.00 – NaN | 0.00 | 0.00 | 1.000 |
|  | Soak time | 3–6 h | 0.27 | 0.08 – 7.13 | 2.74 | 0.87 | 0.382 |
|  | Soak time | 4–5 d | 1.30 | 2.21 – 128 | 12.98 | 2.47 | **0.013** |
|  | Soak time | 8 d | 0.68 | 0.33 – 63.4 | 6.78 | 1.43 | 0.153 |
|  | Soak time | 12–14 d | 0.44 | 0.20 – 19.1 | 4.44 | 1.28 | 0.200 |
|  | Lionfish density |  | 0.10 | 0.10 – 0.11 | 1.00 | -0.17 | 0.866 |
| Native species catch | (Intercept) |  | 0.23 | 0.03 – 1.95 | 0.23 | -1.35 | 0.178 |
|  | Number of traps | Paired | 0.06 | 0.00 – 0.05 | 0.26 | -2.26 | **0.024** |
|  | Distance | 15 m | 0.19 | 0.06 – 0.38 | 0.82 | -0.45 | 0.656 |
|  | Distance | 65 m | 0.03 | 0.00 – 0.08 | 0.14 | -1.32 | 0.188 |
|  | Soak time | 3–6 h | 0.38 | 0.06 – 6.76 | 1.66 | 0.42 | 0.678 |
|  | Soak time | 4–5 d | 1.47 | 1.22 – 72.5 | 6.39 | 1.78 | 0.075 |
|  | Soak time | 8 d | 1.15 | 0.53 – 62.5 | 5.02 | 1.33 | 0.184 |

|  |  |  |  |  |  |  |
| --- | --- | --- | --- | --- | --- | --- |
| Soak time | 12–14 d | 1.51 | 1.22 – 79.8 | 6.56 | 1.77 | 0.077 |
| Lionfish density |  | 0.23 | 0.21 – 0.24 | 0.98 | -0.82 | 0.411 |

\*P-values <0.05 are bolded.

**Fig 5.**
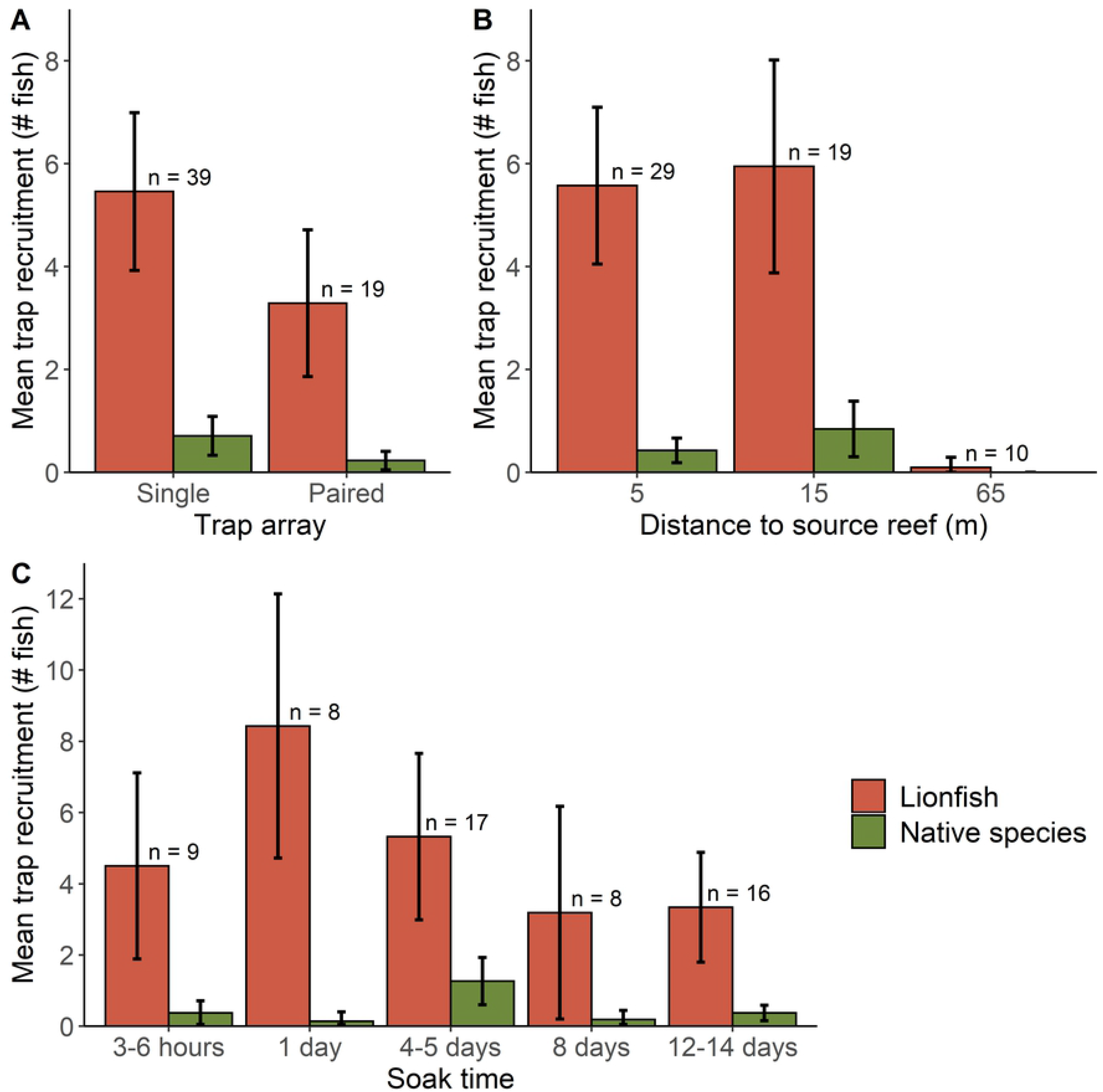
Lionfish and native species recruitment. Mean (± 95 CI) count of lionfish and native species recruitment (fish observed within the trap footprint during retrieval) with number of replicates indicated (n). Trap deployment strategies examined factors of A) distance to source reef, B) trap array, and C) soak time.

Mean native species recruitment and catch were <1 fish per trap (**Fig. 5**). GLMM results indicated native species recruitment and catch were significantly affected by trap number and soak time (**Table 2**). Paired traps had approximately 75% lower recruitment and catch than single traps (**Fig. 5A, Table 2**). In contrast to the lionfish models where recruitment and catch were lower with longer soak times, recruitment and catch for native fishes increased during soak times of 4–5 d, 8 d and 12–14 d (**Fig. 5C**). Longer soak times predicted 4–12X higher recruitment or catch of native species, although given the variance only the 4–5 d level in the native species recruitment model was significant (P = 0.013).

Results from the KS test indicated body sizes (measured in TL) of trap-captured lionfish were significantly larger (P = 0.006) than those of speared fish. Size distributions were multi-modal for both lionfish caught by traps (n = 163) or lionfish speared (n = 3,063) from similar nGOM artificial reefs, likely due to distinct TL modes for juvenile and adult lionfish [51,52]. Mean TL for trap-caught lionfish (277 mm) was 19 mm larger than spear-caught lionfish (296 mm) and showed a right-skewed distribution, indicating traps caught disproportionately fewer juveniles (<200 mm) and more large adult (>300 mm) lionfish (**Fig. 6**). Given a weight-length relationship for nGOM lionfish of *W* = 3.09E-6(*TL*)^3.26 [53], trap-caught lionfish were an average of 73 grams larger.

**Fig 6.**
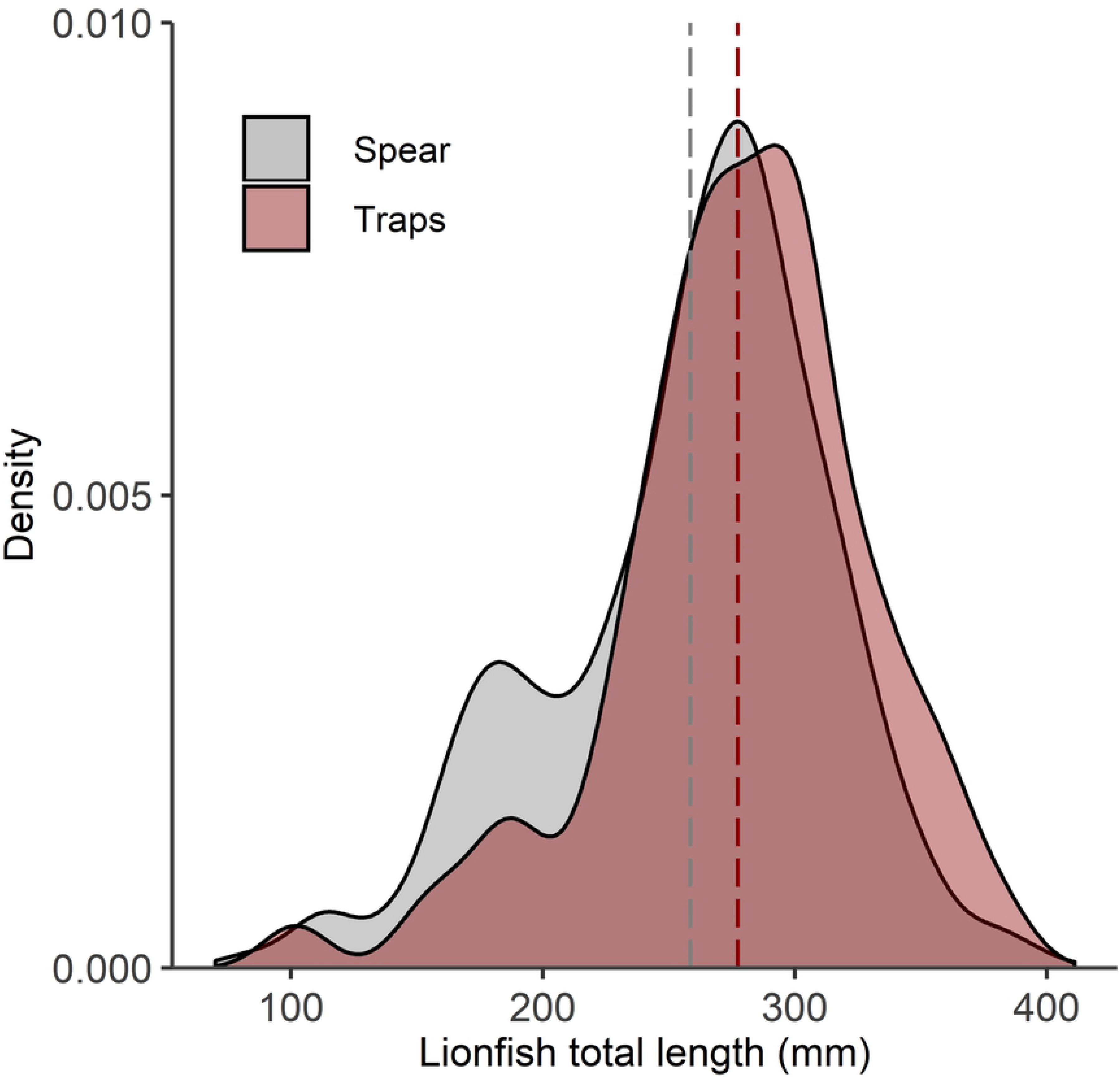
Lionfish size distributions caught by traps or spear. Lionfish total length (TL) distributions for fish captured in Gittings traps (n = 163 lionfish) and spearfishing (n = 3,063 lionfish) with mean TL by gear indicated by vertical lines. Both gear types captured lionfish from northern Gulf of Mexico artificial reefs during June to December 2018.

## DISCUSSION

Testing Gittings traps near nGOM artificial reefs demonstrated an initial proof of concept and identified areas for future work. The lack of apparent environmental impacts indicate traps are suitable for further field testing. Lionfish recruitment and catch were >10X higher than that of native species and traps did not move during severe weather events, although traps will need to be tested for potential trap movement with surface buoys. Lionfish catch was potentially optimized by deploying a single trap 15 m or less from a reef for one day. Mean recruitment with this deployment strategy was approximately 5 lionfish and 0.1 native species per trap. Higher lionfish recruitment to traps deployed closer to reefs would be expected given lionfish exhibit site fidelity [54] and central-place foraging [13]. More sampling at distances between 15 and 65 m would be needed to better understand the effective attraction distance of traps and the linear or non-linear characteristics of that relationship. While variance was high, study results indicated a one-day soak time had the highest lionfish and lowest native species recruitment. This suggests lionfish may recruit relatively quickly to traps, but may leave during longer soak times. The presence of lionfish may also deter native species, as suggested by higher lionfish and lower native species recruitment in shorter soak times and the opposite with longer soak times. Higher lionfish recruitment and catch from single trap deployments disproved our hypothesis that paired deployments could synergistically recruit more lionfish, given lionfish are often found in groups [55] and aggregating behavior is driven by broad-scale habitat complexity [56,57]. Counterintuitively, our results indicated that lionfish density on the adjacent artificial reef was not a significant predictor of lionfish recruitment nor catch. We had expected lionfish movement from reefs to traps would be driven by overcrowding given high lionfish densities have been related to local prey depletion [12,14,58], cannibalism [59,60], and lower body condition [53], and greater movement on coral reefs [61,62].

The current Gittings trap model will need modifications to improve deployment success and decrease lionfish escapement. We suggest design changes include adjustments to flotation and ballast to ensure a vertical orientation during descent to allow successful opening, a reconfigured harness that closes the jaws more quickly and keeps them closed during recovery, a looser net to allow more billowing and not contact lionfish during closure, and faster trap retrieval provided by a winch pulling a buoyed surface line. Additionally, traps for this study were built with relatively thick rebar and the cost to bend the rebar constituted >50% of their construction expense. The thicker rebar caused the traps to weigh >35kg and made moving or deploying the trap by a single person difficult. Either #4 rebar (13 mm / 0.5 in diameter) or #5 rebar (15.875 mm / 0.625” diameter), which are both more easily sourced and manipulated, may provide better materials for trap frames. Concurrent work has examined design and material modifications for reducing production costs of traps (pers. comm., S. Delello, ReefSave) and future testing should test engineering strategies to increase lionfish attraction to traps (e.g., lights, sound, structure). Ultimately, the potential economic viability of a lionfish trap fishery should be carefully considered. Technoeconomic assessments of regional scale deployments should identify capital and operational expenses and evaluate what catch rates could make lionfish trapping commercially feasible.

Removal efficiency by Gittings traps on artificial reefs was 12–26% of the lionfish observed on adjacent artificial reefs. This is higher than many fisheries where removal efficiency is <10% [63–65], but considerably lower than the >85% removal efficiency for spearfishing lionfish on nGOM artificial reefs [46]. Given spearfishing can reduce lionfish densities in areas frequented by divers [66–68] and spearfishing fisheries have caused severe depletion of other reef fishes [69–71], we expect spearfishing to remain the most efficient and cost-effective method for removing lionfish biomass at depths <40 m. Removal efficiencies and uncaptured lionfish will need to be considered when evaluating the potential community benefits offered by lionfish trapping as lionfish removals do not necessarily translate into ecological benefits [14,72,73] given lionfish predation [62,74], growth [53,75], and colonization [14,73,76] rates are controlled via density-dependent feedbacks. Ecosystem models may be appropriate for examining the potential community effects of a deepwater lionfish fishery [15,77].

The selectivity demonstrated by Gittings traps for relatively large lionfish could have implications for depleting their populations. Higher movement rates for larger individuals is reasonable to expect given they are more physically capable to make such movements and face lower risk of predation in transit [78,79]. Predation on lionfish from native species is likely minimal [18,19,80], but juveniles risk conspecific predation [59,81] particularly in areas of high lionfish density [60] where this study was conducted. Smaller lionfish may thus occupy smaller foraging arenas near the reef [82]. Given fecundity increases exponentially with length [83–85], a selectivity bias toward larger fish may have a greater impact on reducing lionfish egg production. High enough reductions would lead to recruitment overfishing whereby the reduced spawning stock decreases future population growth [86]. That said, removal efforts aimed to deplete lionfish biomass should also target juveniles. Harvest of faster-growing juveniles [52,53,87] contributes to growth overfishing [88] and age-structured lionfish population models indicate their exploitation rate is particularly sensitive to lower length at capture [31,32].

The potential for lionfish traps or other novel harvest technologies to reduce lionfish biomass will ultimately be contingent on their capacity to fish them from natural reefs. Natural reefs constitute over 99% of the region’s reef habitat with 90% being mesophotic reefs deeper than 40 m [89]. Even on shallower natural reefs accessible to divers, spearfishing catch rates are limited by lower lionfish density [3,53] and lower removal efficiency [46]. Similar gear testing to that conducted in this study will be needed for traps near natural reefs to evaluate their potential for habitat damage or bycatch, considering there are differences in benthic structure and community composition [14]. It is also currently unknown whether lionfish attraction to traps will be higher near natural versus artificial reefs. Lionfish densities on nGOM artificial reefs are about two orders higher than on nGOM natural reefs [3,46,53], suggesting trap structure may readily attract lionfish from natural reefs. However, if lionfish movement is largely driven by density-dependence or food availability [14,61,62], then the substantially lower densities on natural reefs may limit the effectiveness of lionfish traps near these reefs and the economic viability of a deepwater lionfish fishery.

## Acknowledgements

We thank Josh and Joe Livingston (Dreadknot Charters), Kara Wall (Florida Fish and Wildlife Research Institute), Tony Reyer (NOAA), Sal DeLello (Reef Save), Stacy Frank and Jim Hart (Lionfish University), Laura Tiu (Florida Sea Grant), and Florida Fish and Wildlife Conservation Commission personnel Alan Pierce, Kali Spurgin, Amy Brower, Hanna Tillotson, and Michael Kennison.

## Additional Information

### Author contributions

HH, AF, and SG conceived this study. Sampling design was developed by HH, AF, SG, and RA. HH and AF conducted field operations and data sampling. HH conducted analyses with guidance from RA, MA, and WP. Tables and figures were produced by HH with help from co-authors. HH wrote the manuscript first draft and oversaw revisions with input from co-authors.

### Data availability

All analyses are reproducible with the data and R code provided in the supplementary material.

### Ethics declarations

The authors declare no competing interests. Spearfishers were informed and consented to information about their catch and effort being used for research purposes. Lionfish collection by researchers followed humane sampling protocol with euthanasia via pithing the brain case, as reviewed and approved by the University of Florida’s Institutional Animal Care and Use Committee (UF IACUC Protocol #201810225).

## References

1. Schofield PJ. Update on geographic spread of invasive lionfishes (*Pterois volitans* [Linnaeus, 1758] and *P. miles* [Bennett, 1828]) in the Western North Atlantic Ocean, Caribbean Sea and Gulf of Mexico. Aquat Invasions. 2010;5: 117–122. doi:10.3391/ai.2010.5.S1.024

2. Azzurro E, Stancanelli B, Di Martino V, Bariche M. Range expansion of the common lionfish *Pterois miles* (Bennett, 1828) in the Mediterranean Sea: an unwanted new guest for Italian waters. BioInvasions Rec. 2017;6. doi:10.3391/bir.2017.6.2.01

3. Dahl KA, Patterson WF. Habitat-specific density and diet of rapidly expanding invasive red lionfish, *Pterois volitans*, populations in the northern Gulf of Mexico. PLoS One. 2014;9: e105852. doi:10.1371/journal.pone.0105852

4. Claydon JAB, Calosso MC, Traiger SB. Progression of invasive lionfish in seagrass, mangrove and reef habitats. Mar Ecol Prog Ser. 2012;448: 119–129. doi:10.3354/meps09534

5. Barbour AB, Montgomery ML, Adamson A a., Díaz-Ferguson E, Silliman BR. Mangrove use by the invasive lionfish *Pterois volitans*. Mar Ecol Prog Ser. 2010;401: 291–294. doi:10.3354/meps08373

6. Jud Z, Layman C, Lee J, Arrington D. Recent invasion of a Florida (USA) estuarine system by lionfish *Pterois volitans / P. miles*. Aquat Biol. 2011;13: 21–26. doi:10.3354/ab00351

7. Lesser MP, Slattery M. Phase shift to algal dominated communities at mesophotic depths associated with lionfish (*Pterois volitans*) invasion on a Bahamian coral reef. Biol Invasions. 2011;13: 1855–1868. doi:10.1007/s10530-011-0005-z

8. Andradi-Brown DA, Vermeij MJA, Slattery M, Lesser M, Bejarano I, Appeldoorn R, et al. Large-scale invasion of western Atlantic mesophotic reefs by lionfish potentially undermines culling-based management. Biol Invasions. 2017;19: 939–954. doi:10.1007/s10530-016-1358-0

9. Goodbody-Gringley G, Eddy C, Pitt JM, Chequer AD, Smith SR. Ecological drivers of invasive lionfish (*Pterois volitans* and *Pterois miles*) distribution across mesophotic reefs in Bermuda. Front Mar Sci. 2019;6: 258. doi:10.3389/fmars.2019.00258

10. Gress E, Andradi-Brown DA, Woodall L, Schofield PJ, Stanley K, Rogers AD. Lionfish (*Pterois* spp.) invade the upper-bathyal zone in the western Atlantic. PeerJ. 2017;5: e3683. doi:10.7717/peerj.3683

11. Green SJ, Côté IM. Record densities of Indo-Pacific lionfish on Bahamian coral reefs. Coral Reefs. 2009;28: 107–107. doi:10.1007/s00338-008-0446-8

12. Green SJ, Akins JL, Maljković A, Côté IM. Invasive lionfish drive Atlantic coral reef fish declines. Goldstien SJ, editor. PLoS One. 2012;7: e32596. doi:10.1371/journal.pone.0032596

13. Benkwitt CE. Central-place foraging and ecological effects of an invasive predator across multiple habitats. Ecology. 2016;97: 2729–2739. doi:10.1002/ecy.1477

14. Dahl KA, Patterson III WF, Snyder RA. Experimental assessment of lionfish removals to mitigate reef fish community shifts on northern Gulf of Mexico artificial reefs. Mar Ecol Prog Ser. 2016;558: 207–221. doi:10.3354/meps11898

15. Chagaris D, Binion-Rock S, Bogdanoff A, Dahl K, Granneman J, Harris HE, et al. An ecosystem-based approach to evaluating impacts and management of invasive lionfish. Fisheries. 2017;42: 421–431. doi:10.1080/03632415.2017.1340273

16. Albins MA, Hixon MA. Worst case scenario: Potential long-term effects of invasive predatory lionfish *(Pterois volitans)* on Atlantic and Caribbean coral-reef communities. Environ Biol Fishes. 2013;96: 1151–1157. doi:10.1007/s10641-011-9795-1

17. Hixon MA, Green SJ, Albins MA, Akins JL, Morris JA. Lionfish: a major marine invasion. Mar Ecol Prog Ser. 2016;558: 161–165. doi:10.3354/meps11909

18. Valdivia A, Bruno JF, Cox CE, Hackerott S, Green SJ. Re-examining the relationship between invasive lionfish and native grouper in the Caribbean. PeerJ. 2014;2: e348. doi:10.7717/peerj.348

19. Hackerott S, Valdivia A, Green SJ, Côté IM, Cox CE, Akins L, et al. Native predators do not influence invasion success of Pacific lionfish on Caribbean reefs. Guichard F, editor. PLoS One. 2013;8: e68259. doi:10.1371/journal.pone.0068259

20. Anton A, Simpson MS, Vu I. Environmental and biotic correlates to lionfish invasion success in Bahamian coral reefs. Browman HI, editor. PLoS One. 2014;9: e106229. doi:10.1371/journal.pone.0106229

21. Morris JA, Akins JL, Green SJ, Buddo DSA, Lozano RG. Invasive Lionfish: A Guide to Control and Management. Morris JA, editor. Gulf and Caribbean Fisheries Institute Special Publication Series. Marathon, FL: Gulf and Caribbean Fisheries Institute, Inc.; 2012.

22. Sutherland WJ, Barnard P, Broad S, Clout M, Connor B, Côté IM, et al. A 2017 horizon scan of emerging issues for global conservation and biological diversity. Trends in Ecology and Evolution. Elsevier Ltd; 2017. pp. 31–40. doi:10.1016/j.tree.2016.11.005

23. Johnston MA, Gittings SR, Morris JA. NOAA national marine sanctuaries lionfish response plan (2015-2018): responding, controlling, and adapting to an active marine invasion. 2015. Available: https://repository.library.noaa.gov/view/noaa/13954

24. GEBCO. GEBCO 2019 Grid. 2019. doi:doi:10.5285/836f016a-33be-6ddc-e053-6c86abc0788e

25. Nuttall M, Johnston MA, Eckert RJ, Embesi JA, Hickerson EL, Schmahl GP. Lionfish (*Pterois volitans* [Linnaeus, 1758] and *P. miles* [Bennett, 1828]) records within mesophotic depth ranges on natural banks in the Northwestern Gulf of Mexico. BioInvasions Rec. 2014;3: 111–115. doi:10.3391/bir.2014.3.2.09

26. Switzer TS, Tremain DM, Keenan SF, Stafford CJ, Parks SL, McMichael RH. Temporal and spatial dynamics of the lionfish invasion in the eastern Gulf of Mexico: perspectives from a broadscale trawl survey. Mar Coast Fish. 2015;7: 1–8. doi:10.1080/19425120.2014.987888

27. Reed J, Farrington S, Harter S, Moe H, Hanisak D, David A. Characterization of the mesophotic benthic habitat and fish assemblages from ROV dives on Pulley Ridge and Tortugas during 2014 R/V Walton Smith cruise. 2015. Available: https://repository.library.noaa.gov/view/noaa/17810

28. Andradi-Brown DA, Grey R, Hendrix A, Hitchner D, Hunt CL, Gress E, et al. Depth-dependent effects of culling—do mesophotic lionfish populations undermine current management? R Soc Open Sci. 2017;4: 170027. doi:10.1098/rsos.170027

29. Johnston M, Purkis S. A coordinated and sustained international strategy is required to turn the tide on the Atlantic lionfish invasion. Mar Ecol Prog Ser. 2015;533: 219–235. doi:10.3354/meps11399

30. Johnston MW, Purkis SJ. Hurricanes accelerated the Florida-Bahamas lionfish invasion. Glob Chang Biol. 2015;21: 2249–2260. doi:10.1111/gcb.12874

31. Barbour AB, Allen MS, Frazer TK, Sherman KD. Evaluating the potential efficacy of invasive lionfish (*Pterois volitans*) removals. PLoS One. 2011;6. doi:10.1371/journal.pone.0019666

32. Morris JA, Shertzer KW, Rice JA. A stage-based matrix population model of invasive lionfish with implications for control. Biol Invasions. 2011;13: 7–12. doi:10.1007/s10530-010-9786-8

33. Pitt J, Trott T. Efforts to develop a lionfish-specific trap for use in Bermuda waters. Proceedings of the 66th Gulf and Caribbean Fisheries Institute. Corpus Cristi, TX; 2013. pp. 187–190.

34. Wang Y. Remote operated vehicle for selectively harvesting target species. United States; US10392085B2, 2018.

35. Gittings SR. Apparatus for harvesting lionfish. United States; US20190021297A1, 2018. Available: https://patents.google.com/patent/US20190021297A1/en

36. Gittings SR. Encouraging results in tests of new lionfish trap design. 69th Gulf and Caribbean Fisheries Institute. 2016. pp. 206–214.

37. Gittings SR, Fogg AQ, Frank S, Hart J V, Clark A, Clark B, et al. Going deep for lionfish: designs for two new traps for capturing lionfish in deep water. Silver Spring, MD; 2017. Available: http://www.sanctuaries.noaa.gov

38. Bogdanoff AK, Akins JL, Morris JA, Workgroup 2013 GCFI Lionfish. Invasive lionfish in the marketplace: challenges and opportunities. Proceedings of the 66th Gulf and Caribbean Fisheries Institute. 2014. pp. 140–147.

39. Blakeway RD, Jones GA, Boekhoudt B. Controlling lionfishes (*Pterois* spp.) with consumption: survey data from Aruba demonstrate acceptance of non-native lionfishes on the menu and in seafood markets. Fish Manag Ecol. 2019; fme.12404. doi:10.1111/fme.12404

40. Chapman JK, Anderson LG, Gough CLA, Harris AR. Working up an appetite for lionfish: a market-based approach to manage the invasion of *Pterois volitans* in Belize. Mar Policy. 2016;73: 256–262. doi:10.1016/J.MARPOL.2016.07.023

41. McClanahan TR, Mangi SC. Gear-based management of a tropical artisanal fishery based on species selectivity and capture size. Fish Manag Ecol. 2004;11: 51–60. doi:10.1111/j.1365-2400.2004.00358.x

42. Loh TL, McMurray SE, Henkel TP, Vicente J, Pawlik JR. Indirect effects of overfishing on Caribbean reefs: sponges overgrow reef-building corals. PeerJ. 2015;2015: e901. doi:10.7717/peerj.901

43. Lamberte A, Swingle WE. Amendments to the Reef Fish Fishery Management Plan for the Reef Fish Resources of the Gulf of Mexico. 1993. doi:10.25923/XGHH-CR28

44. Collier C, Cheuvront B, Blough H, Waugh G, McGovern J, MacLauchlin K, et al. Regulatory Amendment 16: Changes to the Seasonal Closure for the Black Sea Bass Pot Sector. 2016. doi:10.25923/2WEY-PQ14

45. Patterson III WF, Dance MA, Addis DT. Development of a remotely operated vehicle based methodology to estimate fish community structure at artificial reef sites in the northern Gulf of Mexico. Proceedings of the 61st Gulf and Caribbean Fisheries Institute. 2009. pp. 263–270.

46. Harris HE, Patterson III WF, Ahrens RNM, Allen MS. Detection and removal efficiency of invasive lionfish in the northern Gulf of Mexico. Fish Res. 2019;213: 22–32. doi:10.1016/j.fishres.2019.01.002

47. Bolker BM, Brooks ME, Clark CJ, Geange SW, Poulsen JR, Stevens MHH, et al. Generalized linear mixed models: a practical guide for ecology and evolution. Trends Ecol Evol. 2009;24: 127–135. doi:10.1016/J.TREE.2008.10.008

48. Bates D, Mächler M, Bolker B, Walker S. Fitting linear mixed-effects models using lme4. J Stat Softw. 2015. doi:10.18637/jss.v067.i01

49. Venables WN, Ripley BD. Modern Applied Statistics with S. 4th ed. New York: Springer; 2002 Available: http://www.stats.ox.ac.uk/pub/MASS4

50. Bolker BM. Ecological Models and Data in R. Princeton University Press; 2007. doi:10.1086/644667

51. Johnson EG, Swenarton MK. Age, growth and population structure of invasive lionfish (*Pterois volitans/miles*) in northeast Florida using a length-based, age-structured population model. PeerJ. 2016;4: e2730. doi:10.7717/peerj.2730

52. Fogg AQ, Ingram W, Peterson MS, Brown-Peterson NJ. Comparing age and growth patterns of invasive lionfish among three ecoregions of the northern Gulf of Mexico. 2015 [cited 12 Mar 2018]. doi:10.13140/RG.2.1.1425.3525

53. Dahl K, Edwards M, Patterson III WF. Density-dependent condition and growth of invasive lionfish in the northern Gulf of Mexico. Mar Ecol Prog Ser. 2019;623: 145–159. doi:10.3354/meps13028

54. Jud ZR, Layman C a. Site fidelity and movement patterns of invasive lionfish, Pterois spp., in a Florida estuary. J Exp Mar Bio Ecol. 2012;414–415: 69–74. doi:10.1016/j.jembe.2012.01.015

55. García-Rivas MDC, Machkour-M’Rabet S, Pérez-Lachaud G, Schmitter-Soto JJ, Doneys C, St-Jean N, et al. What are the characteristics of lionfish and other fishes that influence their association in diurnal refuges? Mar Biol Res. 2017;13: 899–908. doi:10.1080/17451000.2017.1314496

56. Hunt CL, Kelly GR, Windmill H, Curtis-Quick J, Conlon H, Bodmer MD V., et al. Aggregating behaviour in invasive Caribbean lionfish is driven by habitat complexity. Sci Rep. 2019;9. doi:10.1038/s41598-018-37459-w

57. Davis A. Integrating remote sensing and diver observations to predict the distribution of invasive lionfish on Bahamian coral reefs. Mar Ecol Prog Ser. 2019;623: 1–11. doi:10.3354/meps13067

58. Ingeman KE. Lionfish cause increased mortality rates and drive local extirpation of native prey. Mar Ecol Prog Ser. 2016;558. doi:10.3354/meps11821

59. Dahl KA, Patterson III WF, Robertson A, Ortmann AC. DNA barcoding significantly improves resolution of invasive lionfish diet in the northern Gulf of Mexico. Biol Invasions. 2017;19: 1917–1933. doi:10.1007/s10530-017-1407-3

60. Dahl KA, Portnoy DS, Hogan JD, Johnson JE, Gold JR, Patterson WF. Genotyping confirms significant cannibalism in northern Gulf of Mexico invasive red lionfish, *Pterois volitans*. Biol Invasions. 2018;20: 3513–3526. doi:10.1007/s10530-018-1791-3

61. Tamburello N, Côté IM. Movement ecology of Indo-Pacific lionfish on Caribbean coral reefs and its implications for invasion dynamics. Biol Invasions. 2015;17: 1639–1653. doi:10.1007/s10530-014-0822-y

62. Benkwitt C. Invasive lionfish increase activity and foraging movements at greater local densities. Mar Ecol Prog Ser. 2016;558: 255–266. doi:10.3354/meps11760

63. Askey PJ, Richards SA, Post JR, Parkinson EA. Linking angling catch rates and fish learning under catch-and-release regulations. North Am J Fish Manag. 2006;26: 1020–1029. doi:10.1577/M06-035.1

64. Salthaug A, Aanes S. Catchability and the spatial distribution of fishing vessels. Can J Fish Aquat Sci. 2003;60: 259–268. doi:10.1139/f03-018

65. Hangsleben MA, Allen MS, Gwinn DC. Evaluation of electrofishing catch per unit effort for indexing fish abundance in Florida lakes. Trans Am Fish Soc. 2013;142: 247–256. doi:10.1080/00028487.2012.730106

66. De León R, Vane K, Bertuol P, Chamberland VC. Effectiveness of lionfish removal efforts in the southern Caribbean. Endanger Species Res. 2013;22: 175–182. doi:10.3354/esr00542

67. Frazer TK, Jacoby CA, Edwards MA, Barry SC, Manfrino CM. Coping with the lionfish invasion: can targeted removals yield beneficial effects? Rev Fish Sci. 2012;20: 185–191. doi:10.1080/10641262.2012.700655

68. Usseglio P, Selwyn JD, Downey-Wall AM, Hogan JD. Effectiveness of removals of the invasive lionfish: how many dives are needed to deplete a reef? PeerJ. 2017;5: e3043. doi:10.7717/peerj.3043

69. Giglio VJ, Bender MG, Zapelini C, Ferreira CEL. The end of the line? Rapid depletion of a large-sized grouper through spearfishing in a subtropical marginal reef. Perspect Ecol Conserv. 2017;15: 115–118. doi:10.1016/j.pecon.2017.03.006

70. Frisch AJ, Cole AJ, Hobbs J-PA, Rizzari JR, Munkres KP. Effects of spearfishing on reef fish populations in a multi-use conservation area. Ferse SCA, editor. PLoS One. 2012;7: e51938. doi:10.1371/journal.pone.0051938

71. Godoy N, Gelcich S, Vasquez J a., Castilla JC. Spearfishing to depletion: evidence from temperate reef fishes in Chile. Ecol Appl. 2010;20: 1504–1511. doi:10.1890/09-1806.1

72. Harms-Tuohy C, Appeldoorn R, Craig M. The effectiveness of small-scale lionfish removals as a management strategy: effort, impacts and the response of native prey and piscivores. Manag Biol Invasions. 2018;9: 149–162. doi:10.3391/mbi.2018.9.2.08

73. Smith NS, Green SJ, Akins JL, Miller S, Côté IM. Density-dependent colonization and natural disturbance limit the effectiveness of invasive lionfish culling efforts. Biol Invasions. 2017;19: 2385–2399. doi:10.1007/s10530-017-1449-6

74. Benkwitt CE. Non-linear effects of invasive lionfish density on native coral-reef fish communities. Biol Invasions. 2015;17: 1383–1395. doi:10.1007/s10530-014-0801-3

75. Benkwitt CE. Density-dependent growth in invasive lionfish (*Pterois volitans*). Steinke D, editor. PLoS One. 2013;8: e66995. doi:10.1371/journal.pone.0066995

76. Harris HE, Fogg AQ, Allen MS, Ahrens RNM, Patterson WF. Precipitous declines in northern Gulf of Mexico invasive lionfish populations following the emergence of an ulcerative skin disease. Sci Rep. 2020;10: 1–17. doi:10.1038/s41598-020-58886-8

77. Arias-González JE, González-Gándara C, Luis Cabrera J, Christensen V. Predicted impact of the invasive lionfish Pterois volitans on the food web of a Caribbean coral reef. Environ Res. 2011;111: 917–925. doi:10.1016/j.envres.2011.07.008

78. Jenkins DG, Brescacin CR, Duxbury C V., Elliott JA, Evans JA, Grablow KR, et al. Does size matter for dispersal distance? Glob Ecol Biogeogr. 2007;16: 415–425. doi:10.1111/j.1466-8238.2007.00312.x

79. Lima SL, Dill LM. Behavioral decisions made under the risk of predation: a review and prospectus. Can J Zool. 1990;68: 619–640. doi:10.1139/z90-092

80. Hackerott S, Valdivia A, Cox CE, Silbiger NJ, Bruno JF. Invasive lionfish had no measurable effect on prey fish community structure across the Belizean Barrier Reef. PeerJ. 2017;5: e3270. doi:10.7717/peerj.3270

81. Côté IM, Green SJ, Morris JA, Akins JL, Steinke D. Diet richness of invasive Indo-Pacific lionfish revealed by DNA barcoding. Mar Ecol Prog Ser. 2013;472: 249–256. doi:10.3354/meps09992

82. Ahrens RNM, Walters CJ, Christensen V. Foraging arena theory. Fish Fish. 2012;13: 41–59. doi:10.1111/j.1467-2979.2011.00432.x

83. Fogg A, Brown-Peterson N, Peterson M. Reproductive life history characteristics of invasive red lionfish (*Pterois volitans*) in the northern Gulf of Mexico. Bull Mar Sci. 2017;93: 791–813. doi:10.5343/bms.2016.1095

84. Gardner PG, Frazer TK, Jacoby CA, Yanong RPE. Reproductive biology of invasive lionfish (*Pterois* spp.). Front Mar Sci. 2015;2: 7. doi:10.3389/fmars.2015.00007

85. Denney NH, Jennings S, Reynolds JD. Life–history correlates of maximum population growth rates in marine fishes. Proc R Soc London Ser B Biol Sci. 2002;269: 2229–2237. doi:10.1098/rspb.2002.2138

86. Hilborn R, Walters CJ. Quantitative Fisheries Stock Assessment: Choice, Dynamics and Uncertainty. Springer US; 1992. doi:10.1007/978-1-4615-3598-0

87. Fogg AQ, Evans JT, Peterson MS, Brown-Peterson NJ, Hoffmayer ER, Ingram Jr. GW. Comparison of age and growth parameters of invasive red lionfish (*Pterois volitans*) across the northern Gulf of Mexico. Fish Bull. 2019;117: 1–15. doi:10.7755/FB.117.3.1

88. Allen MS, Ahrens RNM, Hansen MJ, Arlinghaus R. Dynamic angling effort influences the value of minimum-length limits to prevent recruitment overfishing. Fish Manag Ecol. 2013;20: 247–257. doi:10.1111/j.1365-2400.2012.00871.x

89. Buczkowski BJ, Reid JA, Jenkins CJ, Reid JM, Williams SJ, Flocks JG. usSEABED: Gulf of Mexico and Caribbean (Puerto Rico and U.S. Virgin Islands) offshore surficial sediment data release. Data Ser. 2006. doi:10.3133/DS146

